# PRL1 and CK2 collaborate to time morning and evening phase in short photoperiods

**DOI:** 10.1101/2022.03.01.482487

**Authors:** Bridget C. Lear, Ela Kula-Eversole, Ravi Allada

## Abstract

The TIMELESS (TIM) phosphatase, *Phosphatase of Regenerating Liver-1 (PRL-1*), is required for setting behavioral phase especially under short photoperiod conditions. Yet the circadian clock neurons and underlying mechanisms important for these effects are unknown. Here we demonstrate that *PRL-1* function in PDF neurons is necessary and sufficient for timing morning and evening phase under short photoperiod (6:18) conditions. In standard (12:12) photoperiod, we find that *PRL-1* rescue or blocking *PRL-1* knockdown in PDF neurons restores wild-type morning phase. Expression of the TIM kinase *shaggy (sgg*) in PDF neurons suppresses in an additive manner *PRL-1* period lengthening and delayed morning phase further underscoring the role of PDF neurons. In contrast to knockdown in all clock neurons, knockdown of *PRL-1* selectively in PDF clock neurons does not alter the timing of morning or evening phase under standard photoperiod conditions, suggesting *PRL-1* function in non-PDF neurons may also be important. In contrast to SGG effects, we find that phase delay of evening onset due to expression of a dominant negative version of the TIM kinase CK2 (CK2alpha^Tik^) requires *PRL-1*. Given that TIM is a substrate of both the PRL-1 phosphatase and CK2 kinase, we hypothesize that PRL-1 dephosphorylation at specific sites may be necessary for subsequent CK2 phosphorylation to time behavior.

## Introduction

The timing of daily behaviors in animals is regulated by networks of clock-containing neurons. The fruit fly *Drosophila* displays crepuscular activity patterns in the presence of daily light:dark cycles, with morning and evening bouts of activity initiated prior to light transitions, as well as sustained ~24 hr rhythms in locomotor behavior in constant darkness. The core molecular clock that drives these daily behaviors involves upregulation of *period (per*) and *timeless (tim*) expression by the transcription factor partners CLOCK (CLK) and CYCLE (CYC), with PER and TIM proteins accumulating at night and translocating to the nucleus to inhibit CLK-CYC activity [1]. Phosphorylation is critical to circadian clock oscillations, and multiple clock proteins exhibit daily rhythms in phosphorylation state. The kinases SHAGGY (SGG, homologous to mammalian GSK3) and CK2 phosphorylate TIM while DOUBLETIME (DBT, homologous to mammalian CK1), CK2, and NEMO phosphorylate PER to regulate nuclear localization, protein stability, and/or protein function [2–10]. Phosphatases have also been implicated in molecular clock function including PHOSPHATASE OF REGENERATING LIVER 1 (PRL-1) which acts on TIM to promote nighttime protein accumulation and proper timing of nuclear translocation [11]. Mutations in or overexpression of core clock components can lead to decreases in rhythmic behavior and/or alterations in behavioral phase and/or period.

The molecular clocks that regulate daily behaviors in *Drosophila* are located in a network of ~150 pacemaker neurons in the brain organized as discrete groups. The large ventral lateral neurons (l-LNv) and four of five small ventral lateral neurons (s-LNv) express the neuropeptide PIGMENT DISPERSING FACTOR (PDF) critical for synchronizing clock oscillations throughout the network and promoting robust free-running rhythmicity [12–14]. PDF-expressing LNv neurons are required for morning activity bouts in anticipation of light transitions and have been designated as ‘morning’ cells (M cells) [15, 16]. Non-PDF expressing neurons including the lateral dorsal neurons (LNd) and a single PDF-negative s-LNv are important for driving evening anticipatory activity during light:dark conditions and are often designated as ‘evening’ cells (E cells) [15, 16]. A subset of dorsal clock neurons (posterior dorsal neurons 1 or DN1p) are coupled to PDF neurons to promote morning behavior, while also contributing to evening behavior depending on environmental conditions [17–19]. The hierarchy of neuronal network effects on free-running rhythmicity as well as morning and evening behavior is dependent on light. M cells determine periodicity in constant darkness and can set the timing of morning and evening behavioral phase in short (winter-like) photoperiod, while E cells are thought to play a more dominant role in regulating rhythmicity and phase in constant light conditions and long photoperiod [16, 20, 21]. The contribution of DN1p to evening behavior is increased in low light or darkness, likely reflecting enhanced coupling to M cells [19]. Studies assessing network hierarchy often utilize period-altering manipulations of clock genes in different circadian neuron subsets. Under standard photoperiods (12 hour light: 12 hour dark), overexpression of period-shortening *sgg* or *Dbt* transgenes in non-PDF clock neurons can advance morning behavior while a period-lengthening *Dbt* transgene can delay morning behavior, consistent with E cell dominance in regulating behavioral timing in standard photoperiod [21, 22]. However, M cell manipulations have mixed effects on morning phase in standard photoperiod, with short period manipulations (*sgg, Dbt^S^*) of PDF neurons sufficient to advanced morning behavior but a long period manipulation (*Dbt^L^*) insufficient to delay morning behavior [21, 22].

We have previously shown that loss of *PRL-1* lengthens circadian period and exhibits phase delaying effects on morning and evening phase especially in short photoperiods. Here we find that the effects of *PRL-1* on morning and evening phase in short photoperiod map to PDF neurons. In addition, we find that *PRL-1* function in both PDF and non-PDF clock neurons contribute to setting phase in standard 12:12 photoperiod. To reveal the underlying mechanism we find that the phase altering effects of the TIM kinase CK2, but not SGG, require *PRL-1* suggesting that PRL-1 dependent TIM phosphorylation sites may be important for CK2 phosphorylation.

## Methods

### *Drosophila* stocks

*Drosophila* strains and crosses were raised and maintained on standard agar food at 22°-25° C in light:dark conditions. *pdfGAL4* [12], *tim*GAL4-62 [23], *pdf*GAL80 [15], *UAS-PRL-1* [24], UAS-*Tik* [25], and UAS-*sgg EP1576* [9], UAS-*sgg^S9E^* [26], *PRL-1* RNAi 45518 [11], *PRL-1^01^* (LL07771-Pbac) [11], and *PRL-1^Df^ (Df(2L) BSC278*) have been described previously. *Df(2L) BSC278* was obtained from Bloomington *Drosophila* stock center (BDSC; Bloomington, IN), *PRL-1(LL07771-Pbac*) was obtained from the *Drosophila* genetics resource center (DGRC; Kyoto), while *PRL-1* RNAi 45518 and UAS-*dcr2* strains were obtained from the Vienna Drosophila Resource Center (VDRC; Vienna).

### Behavioral activity assay and analysis

*Drosophila* locomotor behavioral assays were performed as previously described [11, 27]. Briefly, crosses were raised at 25° C in 12 hour light: 12 hour dark conditions (12L:12D), and ~0-8 day old adult males were loaded into *Drosophila* activity monitors (Trikinetics). To assess *PRL-1* mutant phenotypes, we assayed flies transheterozyous for the protein null allele *PRL-1^01^* and a chromosomal deficiency that removes the *PRL* locus (*Df(2L) BSC278; PRL-1^Df^*) as previously described [11]. Behavioral assays were performed for 5 days of light:dark conditions followed by 7 days constant darkness (DD). For short photoperiod experiments (6 hours light:18 hours dark; 6L:18D), flies were raised at 12L:12D conditions and the lights-off time was advanced by 3-6 H on the day of behavior loading. Group activity profiles and phase and anticipation analysis from 12L:12D and 6L:18D experiments were generated based on normalized, averaged data from the last two days of light:dark conditions to allow for phase adjustment to altered photoperiod regimes [11]. Separate activity profiles were generated from the first day of constant darkness (DD1) after light:dark entrainment. Morning anticipation levels in 12L:12D conditions were determined as previously described [27]. Phase onset timing was determined by first identifying morning and evening peaks of activity from smoothed data profiles, along with the trough values that precede the peak (within 4 hours prior to the morning peak and within 6-8 hours before the evening peak). Phase onset values reflect the time at which 50% of the peak minus trough activity levels are reached. Phase offset measurements for 6L:18D identify peak evening activity and trough activity within 6-8 hours after the peak and again determine the time of 50% activity level between peak and trough. Time windows considered for morning peak activity were as follows: for 12L:12D: *ZT 21-ZT 24;* for DD1 after 12L:12D: *ZT 21-CT4;* for 6L:18D: *ZT17-ZT24;* and for DD1 after 6L:18D: *ZT17-CT1*. Time windows considered for evening peak activity were the following: for 12L:12D: *ZT 6.5 -ZT12*; for DD1 after 12L:12D: *CT 6.5-CT16;* for 6L:18D: *ZT 7-ZT 11.5* (excludes first lights off bin); and for DD1 after 6L:18D: *CT 2- CT12*. Free-running rhythmicity was assessed over 7 days DD using Chi-squared periodogram analysis [27, 28] (Actimetrics). Power measurements indicate chi-squared rhythmic power minus significance (0.01), and rhythmic flies are defined as those with power measurements >= 10. Statistical comparisons were performed using Student’s t-test.

## Results

### *PRL-1* expression in PDF neurons is sufficient to rescue both period and phase delay phenotypes in *PRL-1* mutants especially under short photoperiod

*Drosophila PRL-1* mutants exhibit lengthened free-running period as well as delayed behavioral phase. To map the sites of *PRL-1* function, we assessed period and phase in flies in which *PRL-1* expression is restricted to PDF clock neurons using GAL4-UAS rescue. We find that *pdf*GAL4-driven expression of *UAS-PRL-1* strongly rescues DD period length in *PRL-1* mutants (Table 1, p < .001). Thus, *PRL-1* function in PDF neurons is both necessary and sufficient to regulate free-running period, consistent with a dominant role for M cells in period determination [20]. *PRL-1* mutants also exhibit a loss of morning anticipatory behavior in 12 hour light: 12 hour dark conditions (12L:12D), and a delay in morning and evening behavior on the first day of constant conditions (DD1) after 12L:12D (Fig. 1A,B) [11]. We find that expression of *PRL-1* using *pdfGAL4* significantly restores anticipation in 12L:12D (Fig. 1B,C, top panels) and partial rescue of morning phase on DD1, but no rescue of evening phase onset (Fig. 1B,C, bottom panels). In short photoperiod (6 hours light: 18 hours dark; 6L:18D), *PRL-1* rescue in PDF neurons promotes strong rescue of morning and evening phase phenotypes. *PRL-1* mutants exhibit a ~ 2hr delay in morning phase onset and a ~1H (LD) to ~2.5H (DD1) delay in evening phase offset (Fig. 2A,B). Expression of *PRL-1* in PDF+ neurons rescues both phenotypes with phase measurements comparable to controls (Fig. 2A-C). Taken together these findings support a prominent role for *PRL-1* in PDF+ clock neurons in regulating behavioral phase. However, *PRL-1* expression limited to PDF neurons does not fully restore normal phase in 12L:12D conditions, suggesting *PRL-1* circadian function outside of PDF neurons.

**Figure 1:**
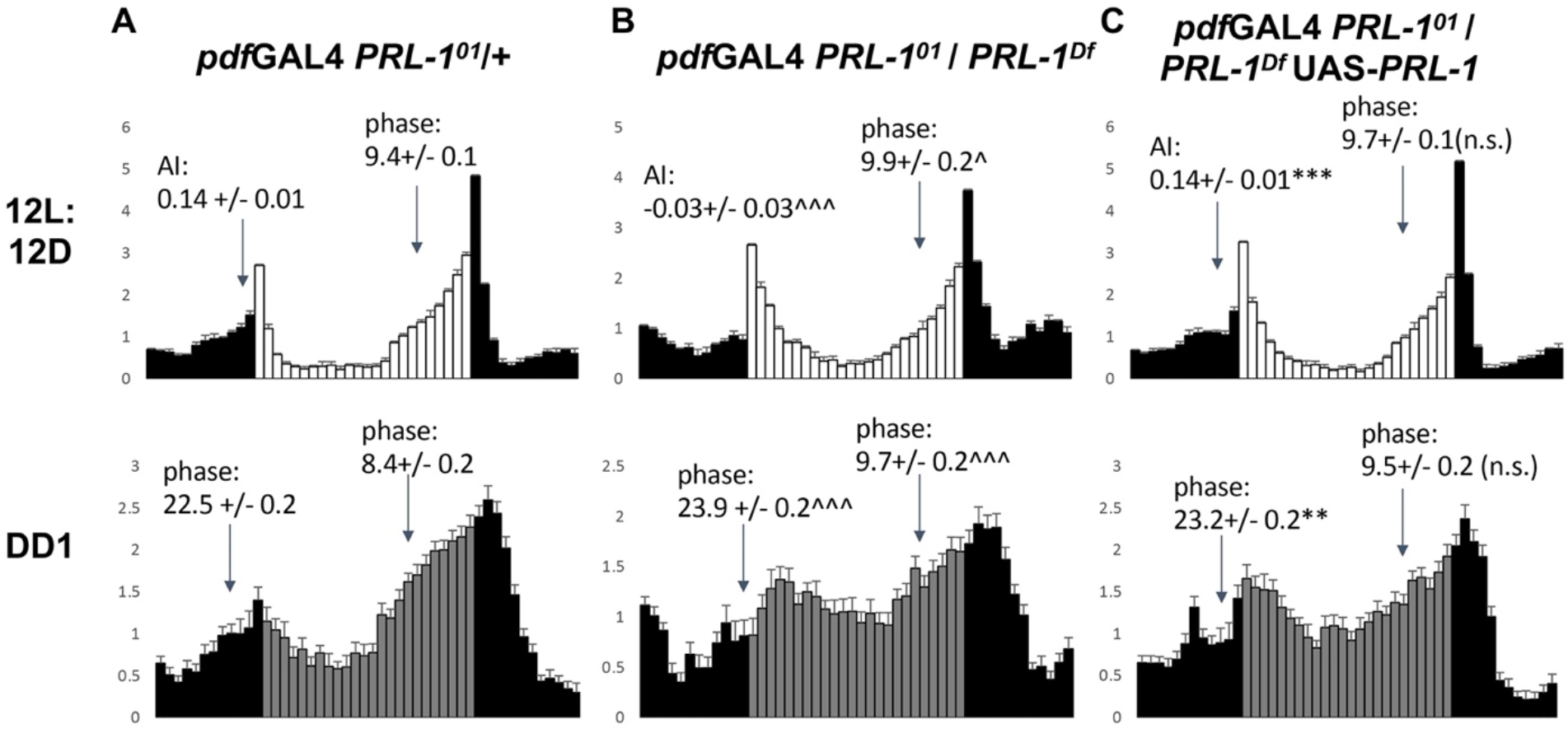
PRL-1 in PDF neurons is sufficient to promote rescue of morning behavior in *Prl-1* mutants in standard photoperiod. Normalized average activity profiles for adult males over the 2 days of 12H Light: 12H Dark (12L:12D) conditions (top panels) and Day 1 of constant darkness (DD1; bottom panels). White bars indicate light phase, gray bars indicate subjective light phase (in DD1), and black bars indicate dark phase. (A) *pdf*GAL4 *PRL-1^01^/+* (n=110) (B) *pdf*GAL4 *PRL-1^0^/PRL-1^Df^* (n=55), and (C) *pdf*GAL4 *PRL-1^0^/PRL-1^Df^ UAS-PRL-1* (n-120). Arrows indicate the time of onset of morning and evening activity bouts, with quantification of phase and anticipation index (for LD morning) indicated (see Methods). Statistical comparisons between control (A) and mutant (B) (^ p < 0.05, ^^^ p < 0.001), and mutant (B) and rescue (C) (** p < 0.01; *** p < 0.001) were performed using Student’s T-test. n.s. = not significant. Error bars indicate SEM.

**Figure 2:**
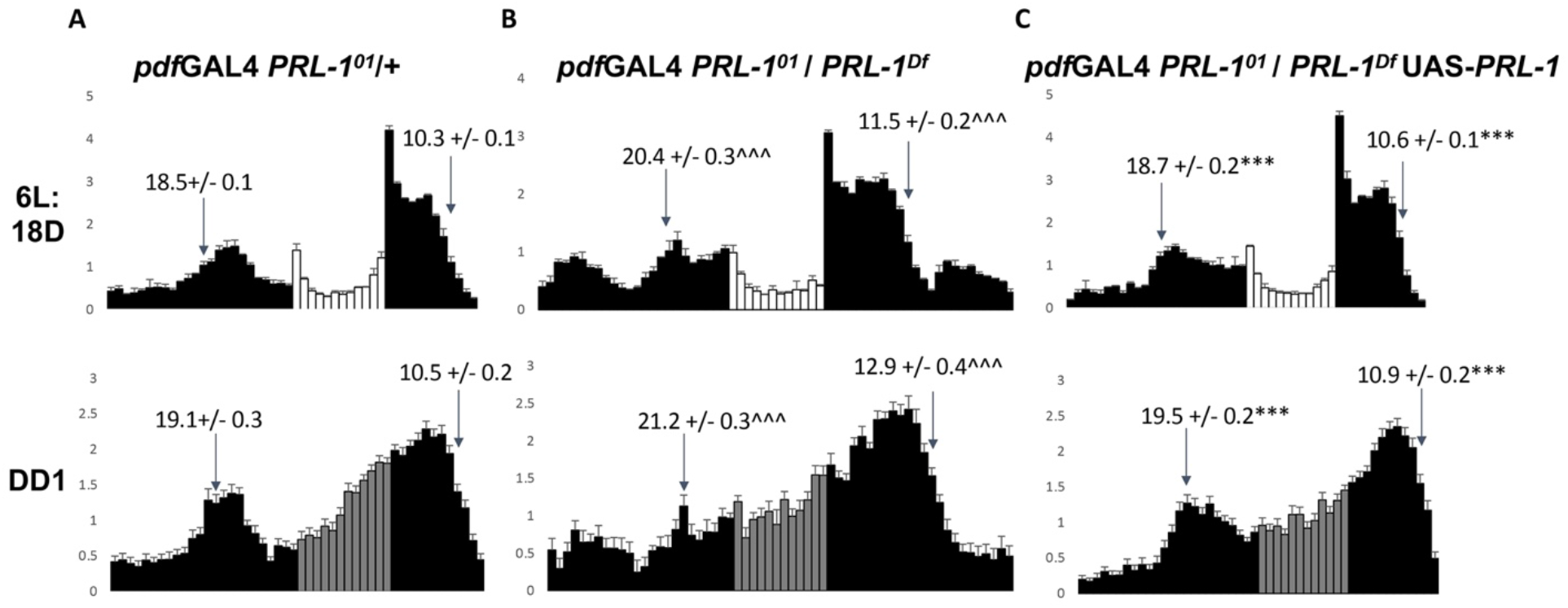
PRL-1 in PDF neurons is sufficient to rescue morning and evening phase delays in *PRL-1* mutants under short photoperiod. Normalized average activity profiles for (A) *pdf*GAL4 *PRL-1^01^*/+ (n=67) (B) *pdf*GAL4 *PRL-1^0^/PRL-1^Df^* (n=38), and (C) *pdf*GAL4 *PRL-1^0^/PRL-1^Df^ UAS-PRL-1* (n=102) over 2 days of 6H Light: 18H Dark (6L:18D) conditions (top panels) and the first day of constant darkness (DD1; bottom panels). White bars indicate light phase, gray bars indicate subjective light phase (in DD1), and black bars indicate dark phase/ subjective dark phase. Arrows indicate the time of onset of morning and evening activity bouts, with quantified average values indicated (see Methods). Significant differences between control (A) and mutant (B) are indicated by ^ (^^^ p < 0.001) and differences between mutant (B) and rescue (C) are indicated by * (*** p < 0.001). Statistical significance determined by Student’s T-test. Error bars indicate SEM.

**Table 1:** PRL-1 expression in PDF clock neurons is sufficient to rescue DD period in *PRL-1* mutants. Analysis of free running locomotor rhythms over 7 days constant darkness (DD) for the genotypes indicated, as determined using chi-squared periodogram. *PRL-1^Df^* refers to *Df(2L) BSC278*, which deletes the *PRL-1* locus. Period and power are measured as average +/− SEM. % Rhythmic indicates percentage of flies for which power measurement (subtracting significance) is >= 10. n indicates number of flies analyzed. Significant differences in period are observed between *PRL-1/+* heterozygotes and *PRL-1^Df^ / PRL-1^01^* transheterozygotes (^^^, p < 0.001) and between *PRL-1* mutants and rescue (***, p < 0.001) as determined using Student’s T-test (p < 0.001).

| Genotype | Period (hrs) | Power | Rhythmic (%) | n |
| --- | --- | --- | --- | --- |
| *PRL-1<sup>Df</sup>/+; UAS-PRL-1/+* | 23.7 +/- 0.1 | 81 +/- 7 | 93 | 40 |
| *pdfGAL4 PRL-1<sup>01</sup>/ +* | 24.3 +/- 0.1 | 81 +/- 4 | 91 | 113 |
| *pdfGAL4 PRL-1<sup>01</sup>/ PRL-1<sup>Df</sup>* | 27.1 +/- 0.1 | 44 +/- 6 | 79 | 34 |
| *pdfGAL4 PRL-1<sup>01</sup>/ PRL-1<sup>Df</sup>; UAS-PRL-1/+* | 24.8 +/- 0.1*** | 55 +/- 4 | 79 | 121 |
| *pdfGAL4 / PRL-1<sup>Df</sup>; UAS-PRL-1/+* | 24.2 +/- 0.2 | 29 +/- 5 | 50 | 40 |

### Requirement for *PRL-1* in timing phase maps to both PDF and non-PDF clock neurons in standard photoperiod

To complement the rescue approach, we also assessed behavioral phenotypes using tissue-specific *PRL-1* RNAi knockdown. Previous findings indicate that *PRL-1* knockdown in all clock neurons (*tim*GAL4) or limited to PDF neurons (*pdf*GAL4) causes robust DD period lengthening comparable to *PRL-1* mutants [11]. In 12L:12D conditions, we observe that *PRL-1* RNAi driven in all or subsets of clock neurons does not promote significant changes in phase or morning anticipation (Fig. 3). However, the timing of light phase in 12L:12D can mask morning phase delays. On DD Day1 after 12L:12D, we observe significant morning and evening phase delays with *PRL-1* knockdown in all clock neurons (Fig. 4A-B; 1.5H morning delay, ~2H evening delay) similar to the delays observed in *PRL-1* mutants. Yet notably, *PRL-1* knockdown limited to PDF neurons does not promote a significant delay in morning or evening phase when compared to both RNAi alone and GAL4 alone controls (Fig. 4A,C). To assess the requirement for *PRL-1* function specifically in non-PDF clock neurons, we drove expression of *PRL-1* RNAi using *tim*GAL4 combined with the GAL4 inhibitor *pdf*GAL80. We have previously found that blocking PDF neuron expression of *PRL-1* RNAi using *tim*GAL4 *pdf*GAL80 suppresses period lengthening [11]. Here we find that restricting *PRL-1* RNAi expression to non-PDF clock neurons also suppresses morning and evening phase delay phenotypes on DD Day1 after 12L:12D (Fig. 4D). This finding differs from other clock manipulations in non-PDF cells, which can advance or delay morning and evening behavior in or after standard photoperiod [21, 22]. In contrast, our data suggest that *PRL-1* mutant 12L:12D phase onset phenotypes reflect a loss of *PRL-1* function in both PDF+ and non-PDF clock neurons. In short photoperiod 6L:18D conditions, RNAi knockdown of *PRL-1* in all clock neurons (Fig. 5A,B) or specifically in PDF neurons (Fig. 5C) promotes significant delays in both morning and evening behavior. On DD Day 1 after 6L:18D, the magnitude of delay phenotypes observed upon PDF neuron knockdown of *PRL-1* is comparable to phenotypes observed upon knockdown in all clock neurons (Fig. 6A-C; ~1.2-1.5H delay in morning, ~1H delay in evening relative to GAL4 only controls). In contrast, *PRL-1* knockdown restricted to non-PDF clock neurons does not promote any detectable phase delay phenotypes in 6L:18D (Fig. 5D) or subsequently on DD Day 1 (Fig. 6D). These findings are consistent with our rescue data, supporting a prominent role of *PRL-1* in PDF neurons in constant darkness or short photoperiod, and a broader anatomical role for *PRL-1* in standard photoperiod.

**Figure 3:**
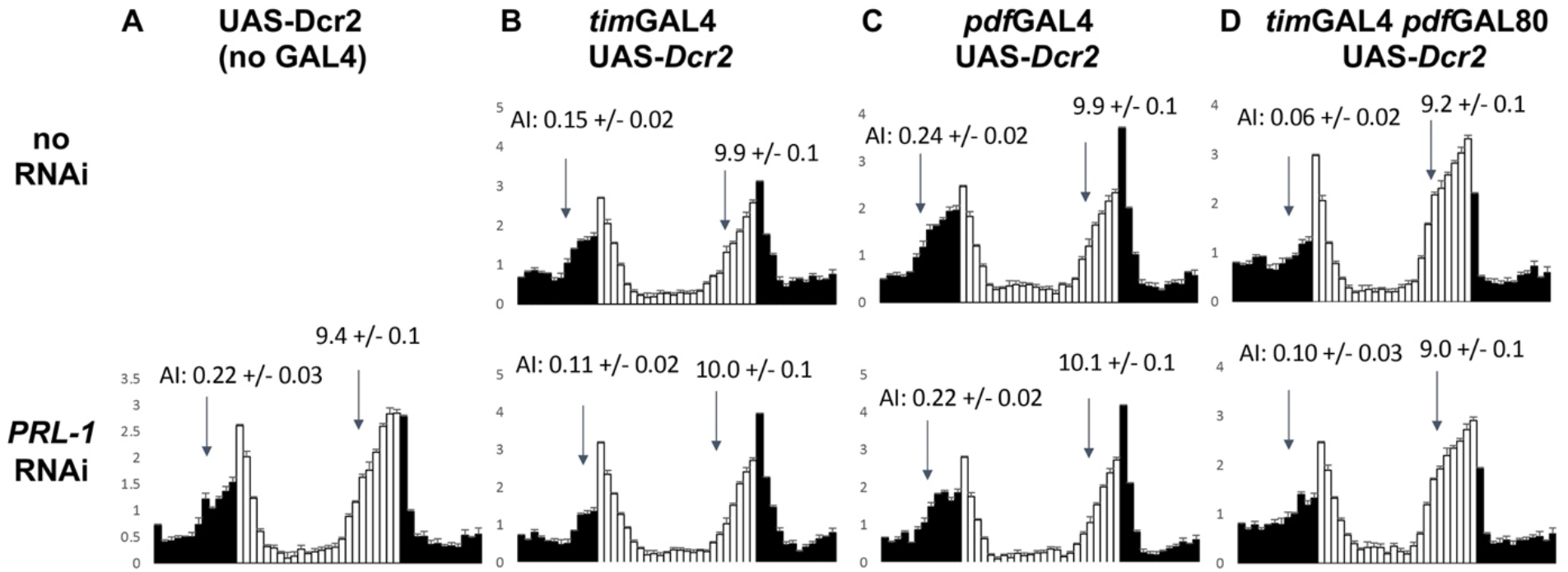
PRL-1 RNAi knockdown in clock neurons does not induce significant changes to M and E behavior in 12:12 light-dark conditions. Normalized average activity profiles for *UAS-Dicer2* (Dcr2) flies combined with (A) no GAL4 (n=28), (B) *timGAL4* (n=55-64) (C) *pdfGAL4* (n=43-45), and (D) *tim*GAL4 *pdf*GAL80 (n=36-38) over 2 days of 12H Light: 12H Dark (LD) conditions. Top panels are groups containing no RNAi and bottom panels are groups containing *UAS-PRL-1* RNAi. White bars indicate light phase and black bars indicate dark phase. Arrows indicate the time of onset of morning and evening activity bouts. For morning behavior, anticipation index (AI) is indicated. For evening behavior, timing of phase onset is indicated (see Methods). No significant phenotypes were detected in GAL4/ RNAi genotypes for either AI or LD phase when compared to both GAL4 alone and RNAi alone, as determined by Student’s T-test. Error bars indicate SEM.

**Figure 4:**
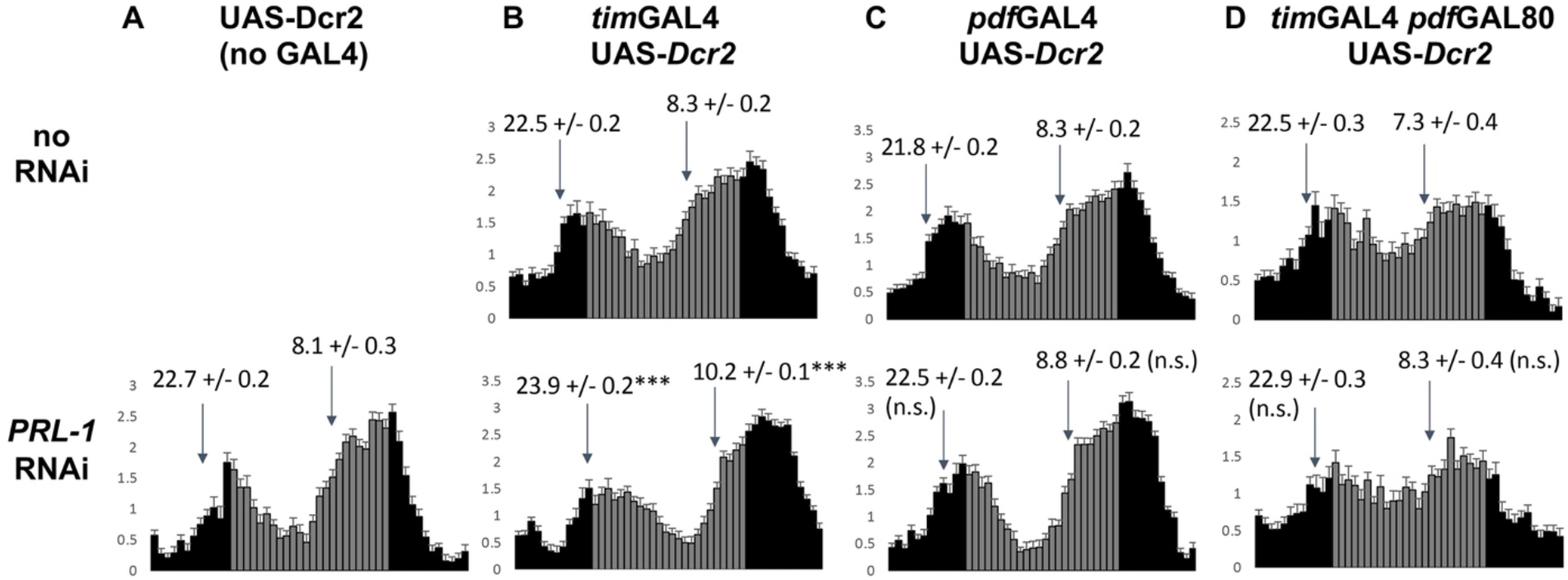
PRL-1 RNAi knockdown in all clock neurons but not in PDF+ or PDF-subsets delays M and E phase after 12L:12D conditions. Normalized average activity profiles for *UAS-Dicer2* (Dcr2) flies combined with (A) no GAL4 (n=28), (B) *tim*GAL4 (n=55-64) (C) *pdf*GAL4 (n=43-45), and (D) *tim*GAL4 *pdf*GAL80 (n=36-38) during DD Day1 after 12H Light: 12H Dark conditions. Top panels are groups containing no RNAi and bottom panels are groups containing *UAS-PRL-1* RNAi. Grey bars indicate subjective day and black bars indicate subjective night. Arrows indicate the time of onset of morning and evening activity bouts, with timing measurement indicated (see Methods). Significant differences between GAL4-RNAi and both controls (GAL4 only, RNAi only) are indicated by * (*** p < 0.001). n.s. indicates not significantly different from both controls. Statistical significance determined by Student’s T-test. Error bars indicate SEM.

**Figure 5:**
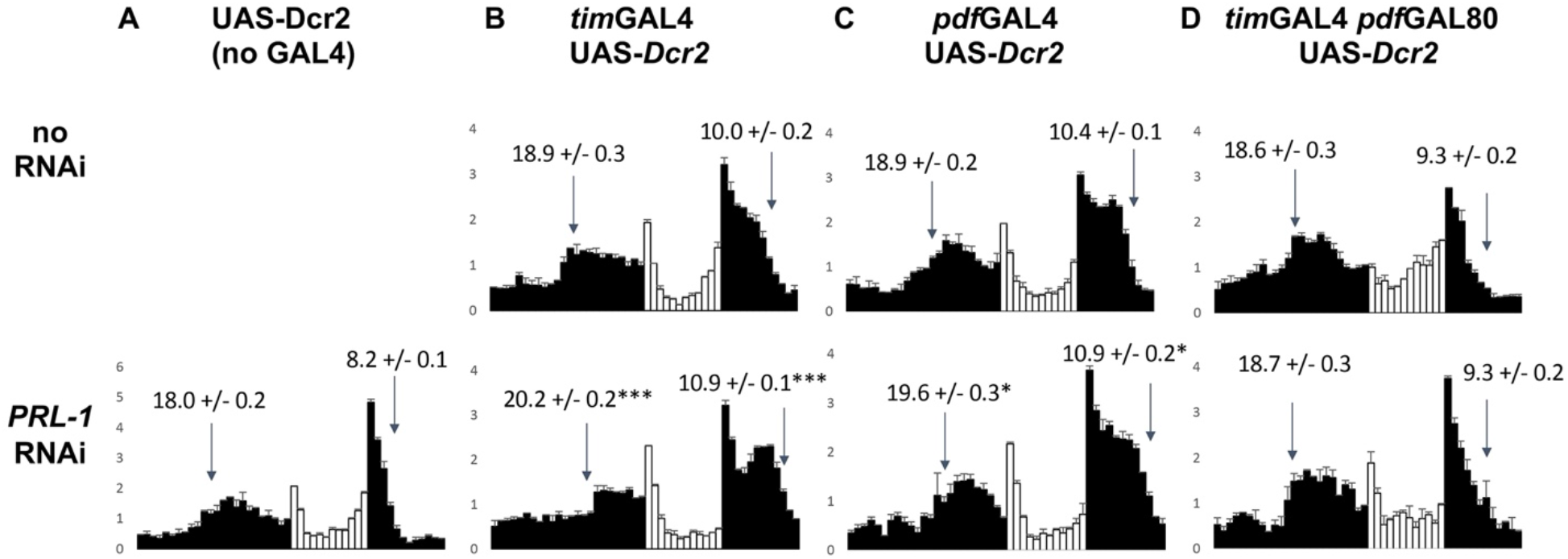
PRL-1 RNAi knockdown in PDF clock neurons delays M and E phase in short photoperiod conditions. Normalized average activity profiles for *UAS-Dicer2* (Dcr2) flies combined with (A) no GAL4 (n=38), (B) *timGAL4* (n=37-92) (C) *pdf*GAL4 (n=33-57), and (D) *timGAL4 pdf*GAL80 (n=21-39) during 2 days of 6H Light: 18H Dark conditions. Top panels are groups containing no RNAi and bottom panels are groups containing *UAS-PRL-1* RNAi. White bars indicate light phase and black bars indicate dark phase. Arrows indicate the time of onset of morning and offset of evening activity bouts, with timing measurement indicated (see Methods). Significant differences between GAL4-RNAi genotypes and both controls (GAL4 only, RNAi only) are indicated by * (* p < 0.05, *** p < 0.001). Statistical significance determined by Student’s T-test. Error bars indicate SEM.

**Figure 6:**
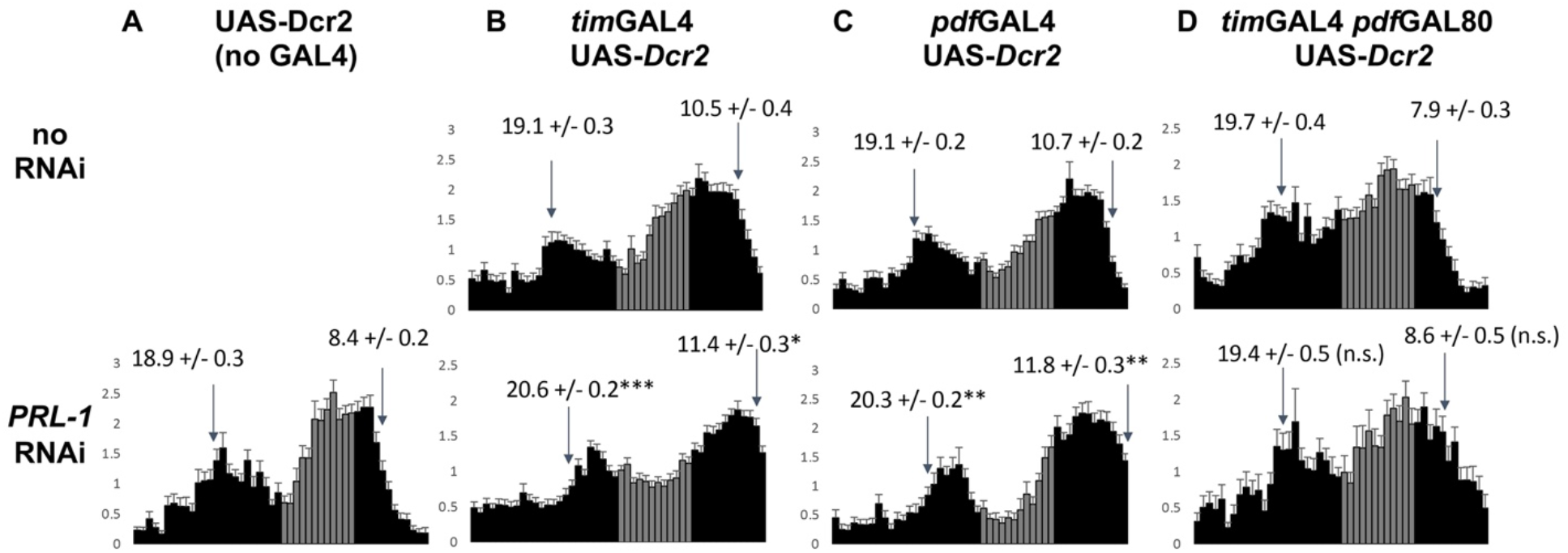
PRL-1 RNAi knockdown in PDF clock neurons delays M and E phase after short photoperiod. Normalized average activity profiles for *UAS-Dicer2* (Dcr2) flies combined with (A) no GAL4 (n=38), (B) *tim*GAL4 (n=37-92) (C) *pdf*GAL4 (n=33-57), and (D) *tim*GAL4 *pdf*GAL80 (n=21-39) during DD Day1 after 6H Light: 18H Dark conditions. Top panels are groups containing no RNAi and bottom panels are groups containing *UAS-PRL-1* RNAi. Grey bars indicate subjective day and black bars indicate subjective night. Arrows indicate the time of onset of morning and offset of evening activity bouts, with timing measurement indicated (see Methods). Significant differences between GAL4-RNAi and both controls (GAL4 only, RNAi only) are indicated by * (* p < 0.05, ** p < 0.01, *** p < 0.001). n.s. indicates not significantly different from both controls. Statistical significance determined by Student’s T-test. Error bars indicate SEM.

### Manipulation of TIM kinases SGG and CK2 exhibit additive effects with *PRL-1* on free-running period

The period-lengthening phenotypes observed in *PRL-1* mutants are accompanied by delayed PER and TIM molecular oscillations in sLNv neurons [11]. To determine the impact of manipulating TIM phosophorylation on *PRL-1* mutant phenotypes, we combined *PRL-1* mutants with transgenic overexpression of the SGG kinase (period shortening) or a dominant-negative CK2alpha kinase allele (*CK2alpha^Tik^*, period lengthening), as well as a genomic *CK2alpha^Tik^* mutant. We find that overexpression of UAS-*sgg* in PDF neurons causes ~ 3 hr period shortening in DD (Table 2) consistent with previous reports [9], whereas *PRL-1^01^/ PRL-1^Df^* mutants exhibit ~3 hr lengthening compared to *pdfGAL4 PRL-1/* + controls (Table 2) [11]. When these two manipulations are combined, we observe an intermediate period phenotype (~24 hr period) that exhibits the expected multiplicative effects of combining two period alterations (13% lengthening of *PRL-1* mutants in control or *sgg* overexpression background) [29]. For the dominant-negative genomic *CK2alpha^Tik^ (Tik*) mutants, we find that combining *Tik/+* heterozygotes (~26 hr period) with *PRL-1* transheterozygous mutants lengthens period to ~28 H (~8%), similar to *PRL-1* mutant lengthening over wild-type (Table 2) [5, 11]. We also used UAS-*Tik* overexpression to restrict CK2 manipulation to PDF neurons, which causes a dramatic period lengthening phenotype (Table 2; 33.5H) [25]. When PDF neuron expression of *UAS-Tik* is combined with loss of *PRL-1*, we also observe near-multiplicative period lengthening (Table 2; ~37H, 10% lengthening over *Tik* alone) despite reduced rhythmicity. Thus, PDF neuron specific manipulations of SGG and CK2 kinases exhibit additive phenotypic effects on free-running period in combination with loss of *PRL-1*, suggesting additive effects on PDF neuron clock speed.

**Table 2:** Additive effects of combining *PRL-1* mutants with *sgg* or *CK2alpha* manipulations on DD period. Analysis of free running locomotor rhythms over 7 days constant darkness (DD) for the genotypes indicated, as determined using chi-squared periodogram. *PRL-1^Df^* refers to *Df(2L) BSC278*, which deletes the *PRL-1* locus. Period and power are measured as average +/− SEM. Rhythmic % indicates percentage of flies for which power measurement (subtracting significance) is >= 10. n indicates number of flies analyzed. Percent (%) lengthening for *PRL-1^Df^/ PRL-1^01^* mutants is calculated relative to the period of the corresponding *pdf*GAL4/ *PRL-1^Df^* strain or *PRL-1^01^/ + Tik/+* strains in the rows directly above the transheterozygous mutant strains.

| Genotype | Period<br>(hrs) | %<br>lengthening | Power | Rhythmic<br>(%) | n |
| --- | --- | --- | --- | --- | --- |
| *pdfGAL4 PRL-1<sup>01</sup>/ +* | 24.2 +/-0.1 |  | 100 +/-6 | 100 | 49 |
| *pdfGAL4 / PRL-1<sup>Df</sup>* | 24.5 +/-0.1 |  | 122 +/-8 | 100 | 25 |
| *pdfGAL4 PRL-1<sup>01</sup>/ PRL-1<sup>Df</sup>* | 27.7 +/-0.2 | 13% | 44 +/-5 | 83 | 42 |
| UAS-*sgg*; *pdfGAL4 / PRL-1<sup>Df</sup>* | 21.2 +/-0.1 |  | 26 +/-3 | 58 | 93 |
| UAS-*sgg*; *pdfGAL4 PRL-1<sup>01</sup>/ PRL-1<sup>Df</sup>* | 24.0 +/-0.1 | 13% | 80 +/-5 | 93 | 98 |
| *pdfGAL4 / PRL-1<sup>Df</sup>*; UAS-*Tik/+* | 33.6 +/-0.2 |  | 35 +/-5 | 82 | 33 |
| *pdfGAL4 PRL-1<sup>01</sup>/ PRL-1<sup>Df</sup>*; UAS-*Tik/+* | 37.0 +/-0.3 | 10% | 16 +/-3 | 42 | 31 |
| UAS-*sgg*; *pdfGAL4 / PRL-1<sup>Df</sup>* | 21.2 +/-0.1 |  | 26 +/-3 | 58 | 93 |
| UAS-*sgg*; *pdfGAL4 PRL-1<sup>01</sup>/ PRL-1<sup>Df</sup>* | 24.0 +/-0.1 | 13% | 80 +/-5 | 93 | 98 |
| *Tik/+* | 26.2 +/-0.0 |  | 111 +/-7 | 90 | 42 |
| *PRL-1<sup>01</sup>/+; Tik/+* | 26.0 +/-0.2 |  | 81 +/-15 | 91 | 11 |
| *PRL-1<sup>01</sup>/ PRL-1<sup>Df</sup>*; *Tik/+* | 27.9 +/-0.1 | 8% | 45 +/-5 | 81 | 48 |

### Overexpression of *sgg* in PDF neurons suppresses *PRL-1* phase delays in an additive manner

To determine how manipulations of TIM phosphorylation impact behavioral phase with *PRL-1*, we assessed LD and DD Day1 behavioral profiles in *PRL-1* mutants combined with PDF-neuron expression of *sgg*. We find that *pdfGAL4* driven expression UAS-*sgg* promotes a modest advance of morning phase in 12L:12D and DD Day1 but no advance of evening behavior, consistent with previous findings (Fig. 7A,C) [21]. In a *PRL-1* mutant background, PDF neuron expression of *sgg* restores morning anticipation in 12L:12D (Fig. 6B,D, top panels) and normal timing of morning phase onset on DD Day 1 but with no significant effect on DD1 evening phase onset (Fig. 7B,D, bottom panels). Thus, *sgg* expression specifically in PDF neurons is sufficient to suppress *PRL-1* morning phase and anticipation phenotypes in 12L:12D conditions. We also assessed the combined effects of *PRL-1* and *sgg* expression on phase in short photoperiod conditions. In these experiments we overexpressed a mutant version of *sgg* (*UAS-sgg^S9E^*) in PDF neurons that promotes advanced evening behavior in 6L:18D but without a consistent or significant advance in morning behavior (Fig. 8A,C) [26]. The combination of *pdf*GAL4-driven expression of *sgg^S9E^* with *PRL-1* mutants in short photoperiod again exhibits an additive effect, with significant delays in evening behavior in *PRL-1* mutants (Figure 9A,B) suppressed by *sgg^S9E^* expression (Figure 9D). Thus expression of *sgg* specifically in PDF neurons suppresses multiple behavioral phenotypes in *PRL-1* mutants including period lengthening and phase delay phenotypes.

**Figure 7:**
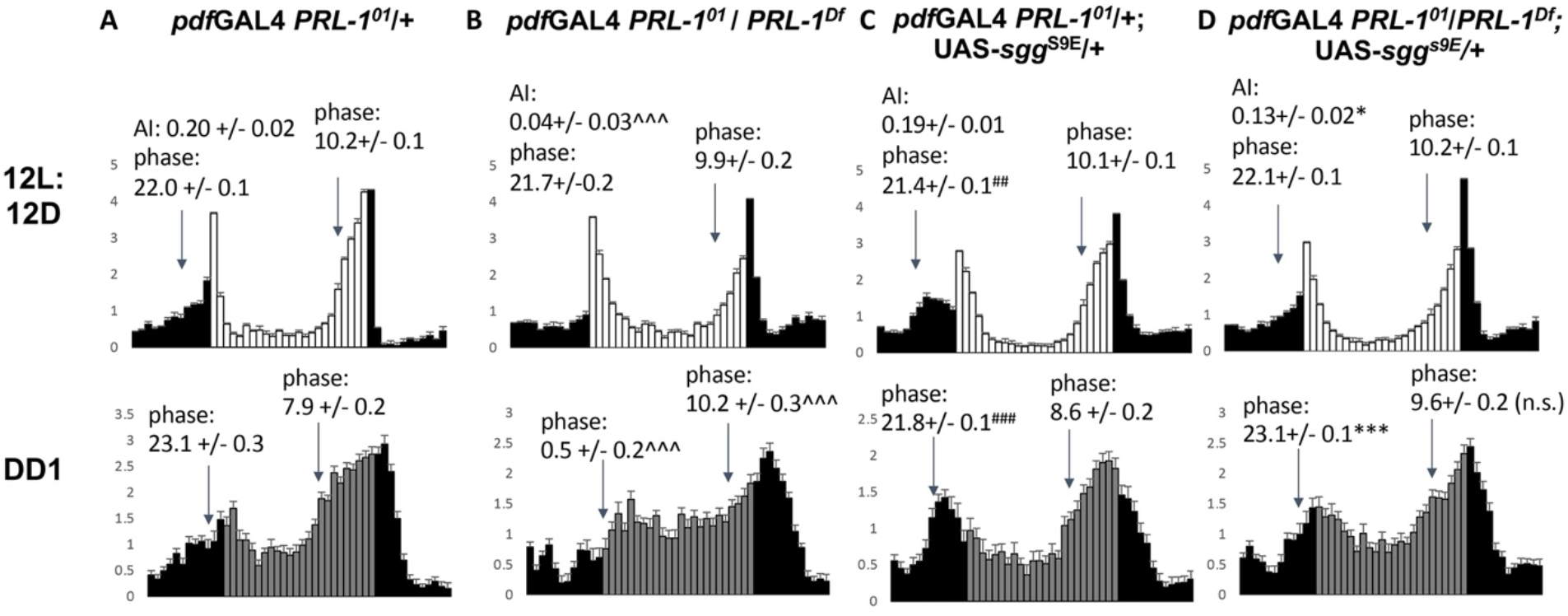
Overexpression of SGG in PDF neurons is sufficient to restore morning anticipation and phase in *Prl-1* mutants in standard photoperiod. Normalized average activity profiles for (A) *pdf*GAL4 *PRL-1^01^*/+ (n=49) (B) *pdf*GAL4 *PRL-1^01^/ PRL-1^Df^* (n=43), and (C) UAS-*sgg pdf*GAL4 *PRL-1^01^*/+ (n=94) (D) UAS-*sgg pdf*GAL4 *PRL-1^01^/ PRL-1^Df^* (n=105) over 2 days of 12H Light: 12H Dark (LD) conditions (top panels) and the first day of constant darkness (DD1; bottom panels). White bars indicate light phase, gray bars indicate subjective light phase (in DD1), and black bars indicate dark phase/ subjective dark phase. Arrows indicate the time of onset of morning and evening activity bouts, with quantified average values indicated (see Methods). Morning anticipation index (AI) is indicated for LD conditions (see Methods). Significant differences between control (A) and mutant (B) are indicated by ^ (^^^ p < 0.001), between control (A) and sgg overexpression (C) are indicated by # (## p < 0.01, ### p < 0.001), and between mutant (B) and mutant plus sgg overexpression (D) are indicated by * (* p < 0.01; *** p < 0.001). n.s. = not significant. Statistical significance determined by Student’s T-test. Error bars indicate SEM.

**Figure 8:**
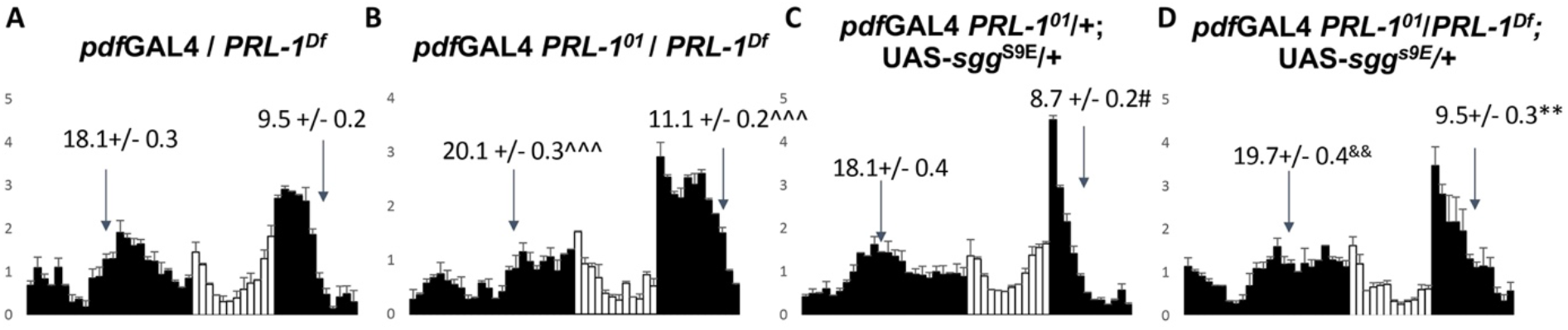
Overexpression of SGG^S9E^ in PDF neurons has additive effects with *PRL-1* in short photoperiod. Normalized average activity profiles for (A) *pdfGAL4/ PRL-1^Df^* (n=9) (B) *pdfGAL4 PRL-1^01^/ PRL-1^Df^* (n=23), and (C) *pdfGAL4 PRL-1^01^/+; UAS-sgg^S9E^/+* (n=47) (D) *pdfGAL4 PRL-1^01^/ PRL-1^Df^*; UAS-*sgg^S9E^*/+ (n=20) over 2 days of 6H Light: 18H Dark (6L:18D) conditions. White bars indicate light phase and black bars indicate dark phase. Arrows indicate the time of onset of morning and evening activity bouts, with quantified average values indicated (see Methods). Significant differences between control (A) and mutant (B) are indicated by ^ (^^^ p < 0.001), and between control (A) and sgg overexpression (C) are indicated by # (# p < 0.05). For sgg overexpression in a mutant background (D), evening phase offset timing is intermediate between mutant (B) and overexpression (C) measurements (** p < 0.01), and morning phase is significantly delayed compared to sgg overexpression alone (C; && p < 0.01). Statistical significance determined by Student’s T-test. Error bars indicate SEM.

**Figure 9:**
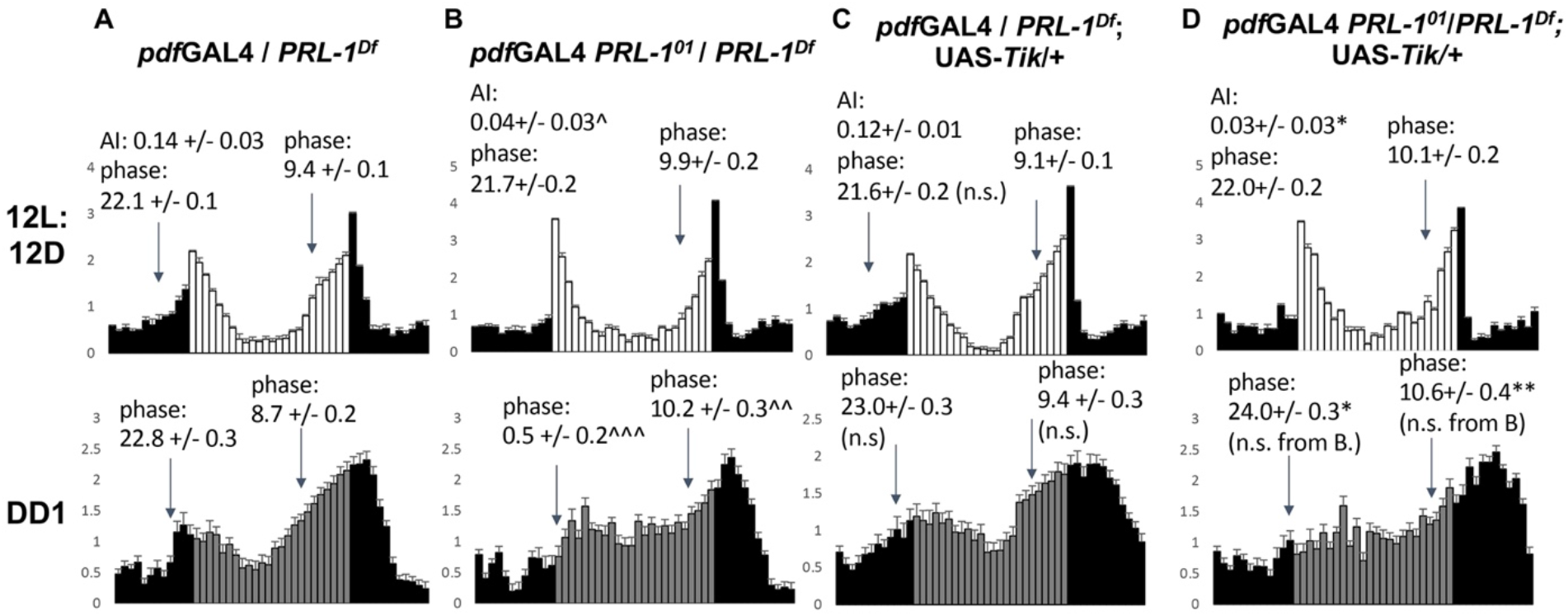
Overexpression of Tik in PDF neurons does not delay phase onset in standard photoperiod. Normalized average activity profiles for (A) *pdf*GAL4/ *PRL-1^Df^* (n=25) (B) *pdf*GAL4 *PRL-1^01^/ PRL-1^Df^* (n=43), and (C) *pdf*GAL4/ *PRL-1^Df^;* UAS-*Tik*/+ (n=35) (D) *pdf*GAL4 *PRL-1^01^/ PRL-1^Df^;* UAS-*Tik*/+ (n=31) over 2 days of 12H Light: 12H Dark (LD) conditions (top panels) and the first day of constant darkness (DD1; bottom panels). White bars indicate light phase, gray bars indicate subjective light phase (in DD1), and black bars indicate dark phase/ subjective dark phase. Arrows indicate the time of onset of morning and evening activity bouts, with quantified average phase values indicated (see Methods). Morning anticipation index (AI) is indicated for LD conditions (see Methods). Significant differences between control (A) and mutant (B) are indicated by ^ (^ p < 0.05, ^^ p < 0.01, ^^^ p < 0.001), and between control (A) and mutant plus Tik expression (D) are indicated by * (* p < 0.05; ** p < 0.01). No significant differences were detected between control (A) and Tik overexpression (C), or between mutant (B) and mutant plus Tik expression (D) for phase onset and morning anticipation. n.s. = not significant. Statistical significance determined by Student’s T-test. Error bars indicate SEM.

### PRL-1 is required for Tik effects on morning and evening phase in short photoperiod

We also tested a period-lengthening manipulation of CK2 by combining *PRL-1* mutants with PDF-neuron specific expression of *UAS-Tik*. In 12L:12D, *pdf*GAL4 expression of UAS-*Tik* has little effect on phase onset despite large effects on period (Fig. 9, A-C). This is similar to our findings with PDF neuron-specific knockdown of *PRL-1* (Fig. 4), which fails to phenocopy the delayed morning and evening phase onset phenotypes of *PRL-1* mutants (Fig. 9, A-B, bottom panels). Expression of *UAS-Tik* with *pdf*GAL4 also does not further delay morning or evening phase onset in *PRL-1* mutants after 12L:12D (Fig. 9, B,D). In short photoperiod conditions, (6L:18D) we find that *pdf*GAL4-driven expression of *UAS-Tik* results in a significant delay in both morning and evening phase onset similar to *PRL-1* single mutants (Fig. 10, A-C). Yet surprisingly, when we combine PDF neuron *Tik* expression with loss of *PRL-1*, behavioral phase is not further delayed when compared to *PRL-1* mutants (Fig. 10,B,D). Evening phase offset actually occurs significantly earlier in *pdf*GAL4 *UAS-Tik* in a *PRL-1* mutant background (10.8 +/− 0.2) compared to *pdf*GAL4 UAS-*Tik* in PRL-1/+ heterozygotes (12.0 +/− 0.1; p < 0.001). These data suggest that the phase delaying effects of *CK2* manipulations require *PRL-1* function in short photoperiod.

**Figure 10:**
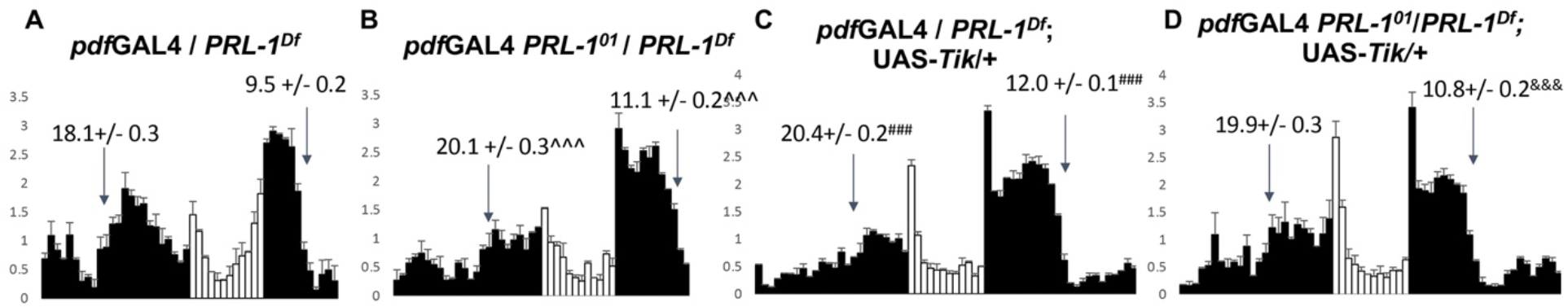
Phase delays induced by PDF neuron expression of CK2alpha-Tik in short photoperiod require *PRL-1*. Normalized average activity profiles for (A) *pdfGAL4/ PRL-1^Df^* (n=9) (B) *pdf*GAL4 *PRL-1^01^/ PRL-1^Df^* (n=23), and (C) *pdf*GAL4 *PRL-1^01^/+;* UAS-*Tik*/+ (n=46) (D) *pdf*GAL4 *PRL-1^01^/ PRL-1^Df^; UAS-Tik/+* (n=28) over 2 days of 6H Light: 18H Dark (6L:18D) conditions. White bars indicate light phase and black bars indicate dark phase. Arrows indicate the time of onset of morning and evening activity bouts, with quantified average values indicated (see Methods). Significant differences between control (A) and mutant (B) are indicated by ^ (^^^ p < 0.001), and between control (A) and Tik overexpression (C) are indicated by # (### p < 0.001). For Tik overexpression in a mutant background (D), morning phase onset and evening phase offset are not significantly different from *PRL-1* mutants (B), while evening phase offset is significantly earlier than Tik overexpression (&&& p < 0.001). Statistical significance determined by Student’s T-test. Error bars indicate SEM.

## Discussion

Here we have identified novel roles for *PRL-1* in regulating behavioral phase. Specifically, we find that *PRL-1* is required for PDF neuron manipulations of *CK2alpha^Tik^* to delay phase under short photoperiod, suggesting functional interactions between PRL-1 phosphatase and CK2 kinase, which both act on TIM. We also find that the requirement for *PRL-1* in phase regulation under standard photoperiod maps to both PDF and non-PDF neurons. These results indicate a unique anatomical role for *PRL-1* in regulating phase in standard photoperiod compared to other clock genes that have been examined.

We find that *PRL-1* is required for the effects of PDF-neuron *Tik* expression on phase under short photoperiod. Both the loss of *PRL-1* and PDF neuron-specific expression of *Tik* cause morning and evening phase delays in short photoperiod, but the combination of these manipulations results in phase profiles that are indistinguishable from *PRL-1* mutants alone. Notably, both PRL-1 phosphatase and CK2 kinase are known to act directly on TIM [5, 8, 10, 11]. Loss of *PRL-1* phosphatase results in delayed nighttime accumulation and nuclear entry of TIM in PDF neurons [11], while decreased CK2 function through *Tik* expression leads to elevated TIM levels and delayed nuclear entry [10, 25]. One potential explanation for our behavioral findings is that TIM dephosphorylation by PRL-1 may be required for subsequent CK2 phosphorylation under short photoperiod conditions. Thus *Tik* expression in PDF neurons would not cause further delays to M cell clocks in the absence of *PRL-1*. Alternatively, the combined effects of *CK2* and *PRL-1* manipulations may delay PDF neuron clock phase to such an extent that the neurons are less effective at delaying clocks in non-PDF neurons to impact phase. Large discrepancies in clock speed between PDF and non-PDF neurons are known to limit synchronization of clocks in the circadian network, causing rhythms to split into two free-running components [30]. Yet here we find that PDF neurons appear to retain some ability to delay morning evening behaviors (~1.5-2 hr relative to controls), suggesting that PDF neurons in *pdfGAL4 UAS-Tik PRL-1* mutant flies can still delay non-PDF clocks to some extent under light:dark entrainment conditions. Clock manipulations that include PDF neurons often produce more modest effects on phase under entrainment conditions when compared to the magnitude of period alterations [21, 22, 25]. However, to our knowledge a limiting effect of combined clock alterations on phase advances or delays has not been described previously. It will be relevant to determine the molecular effects of combining *PRL-1* and *CK2* manipulations under short photoperiod, and to assess whether other period lengthening manipulations show non-additive effects on phase in combination with *PRL-1* mutants and/or PDF neuron expression of *Tik*.

Our data also reveal unexpected anatomical requirements for *PRL-1* in the regulation of behavioral phase. We find that RNAi knockdown of *PRL-1* in all clock neurons can phenocopy the phase delay phenotypes of *PRL-1* mutants in standard photoperiod, but that knockdown in either the PDF or non-PDF subsets does not cause significant delays. Previous findings with both period shortening (*sgg*) and period lengthening (*Dbt^L^*) manipulations indicate that clock manipulations in non-PDF neurons (or more specifically in E cells) in standard photoperiod shifts the timing of both morning and evening behavior [21, 22]. Notably, *PRL-1* is expressed broadly in the clock network, but rhythmic expression has only been detected within the PDF neuron subset [11]. Thus it is possible that loss of *PRL-1* may have different molecular effects in PDF and non-PDF clock neurons. Our findings also differ from previous reports in the effects of M cell manipulations on evening behavior on DD Day1. We find that *PRL-1* knockdown in PDF neurons does not delay evening onset timing, whereas DD1 evening delays have previously been reported with other PDF specific clock manipulations [22, 25, 31]. This may reflect differences in quantification methods, in that our current method assesses evening phase onset while previous studies have evaluated peak and/or activity offset. Network manipulations may differentially impact the onset, peak, and/or offset of activity peaks.

Our results also address the differential effects of M cell clock manipulations on morning behavior under standard photoperiod. Under 12L: 12D conditions we find that PDF neuron specific knockdown of *PRL-1* or expression of UAS-*Tik* does not delay morning phase despite large effects on period. These findings differ from findings with *sgg* overexpression, which advances morning behavior in standard photoperiod [21], but are consistent with previous data for a period-lengthening manipulation *Dbt^L^* [22]. We also show that these same manipulations (*pdf*GAL4 UAS Tik or *pdf*GAL4 *UAS-PRL-1* RNAi) can induce substantial phase delays in short photoperiod, thus the effects are photoperiod-dependent and consistent with M cell dominance in short photoperiod [21]. The differential ability of PDF neurons to delay or advance non-PDF clocks in non-PDF neurons may contribute to the different effects on behavioral phase in standard photoperiod [22], although the clock effects of such manipulations across the circadian network have not been examined on DD Day1.

## Acknowledgements

We would like to thank Miguel del Busto, Esha Kular, Dahyun Lee, and Ima Samba for assistance with behavioral assays. We thank the Bloomington *Drosophila* Stock Center and Vienna *Drosophila* Resource Center for fly stocks. The research was supported by NIH (R01NS106955) to R.A.

## Notes

### Competing Interest Statement

The authors have declared no competing interest.

